## Supplementary Information for "Lipidomic and biophysical homeostasis of mammalian membranes in response to dietary lipids is essential for cellular fitness"

Supplementary Information for Levental et al "**Homeostatic remodeling of mammalian membranes in response to dietary lipid perturbations is essential for cellular fitness**"

Contents: 14 supplementary figures and one supplementary table

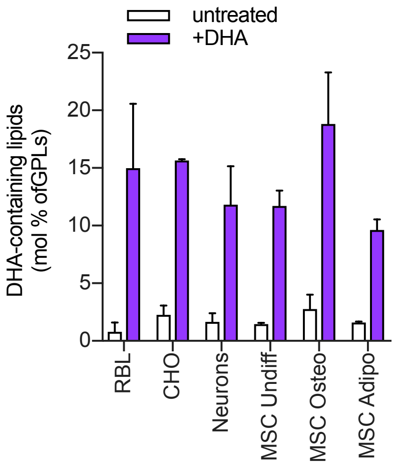


**Fig S1. DHA is robustly incorporated into membranes of various cell types**. mol% of DHA-containing lipids as a fraction of all cellular phospholipids. RBL and CHO cells were treated for 3 days with 20 μM DHA; rat hippocampal neurons were supplemented with 3.5 μM DHA in NS21 medium for 17 days; human MSC were cultured and/or differentiated in the presence of 20 μM DHA for 14 days[1]. Average ± SD of n ≥ 3 independent experiments. The data for all MSC samples was reported in our previous publication[1].

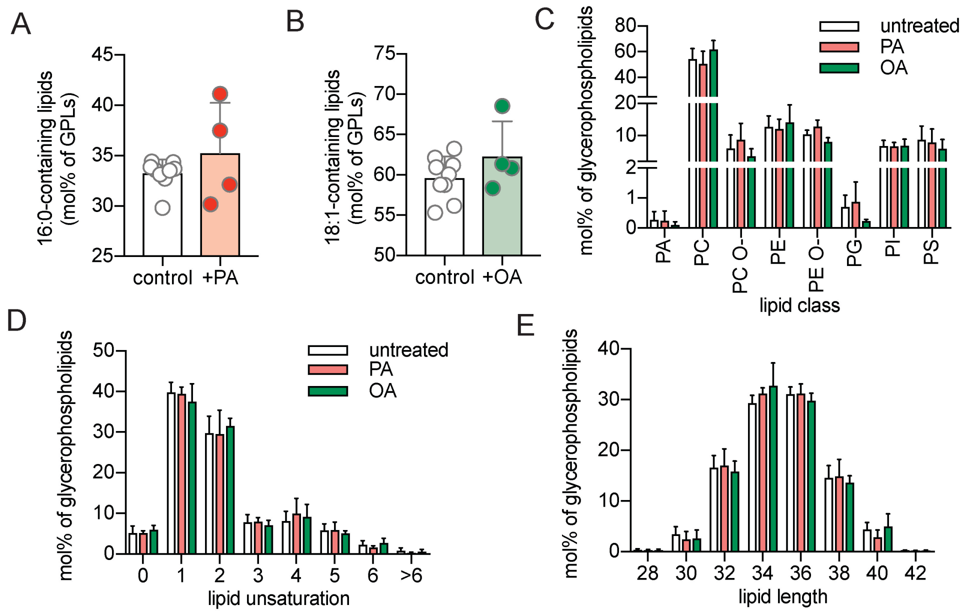


**Fig S2. No notable lipidomic effects of PA or OA supplementation.** Supplementation with (A) 20 μM palmitic acid (PA) and (B) oleic acid (OA) did not result in a significant increase in lipids containing either PA or OA, respectively. Supplementation with either PA or OA has little effect on the cellular lipidomes, consistent with their increased lack of incorporation. No major effects were observed for (C) glycerophospholipids headgroup distribution, (D) lipid unsaturation, and (E) lipid length. Average ± SD of n ≥ 4 independent experiments.

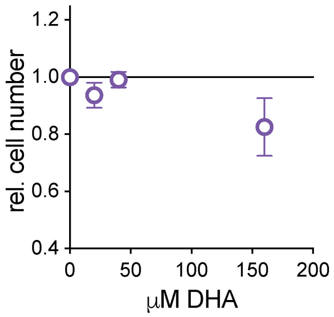


**Fig S3. No toxicity of DHA at lower doses**. At low doses (20-40 µM), DHA has minimal effects on cell health. At higher doses, DHA is slightly cytostatic. All experiments in this manuscript are performed with 20 µM DHA supplementation, which alone causes no effect on cell survival or proliferation.

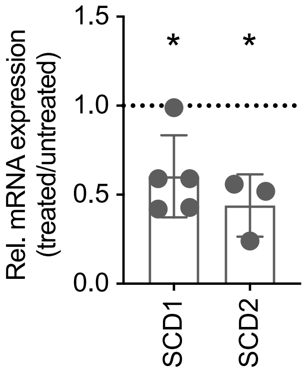


**Fig S4. Transcriptional downregulation of desaturases.** Changes in gene expression of fatty acid desaturases after 24 h of supplementation with DHA compared to unsupplemented cells (measured by qPCR via the delta delta Ct method). Average ± SD of n ≥ 3 independent experiments. *p<0.05 as determined by one-sample t-test to untreated.

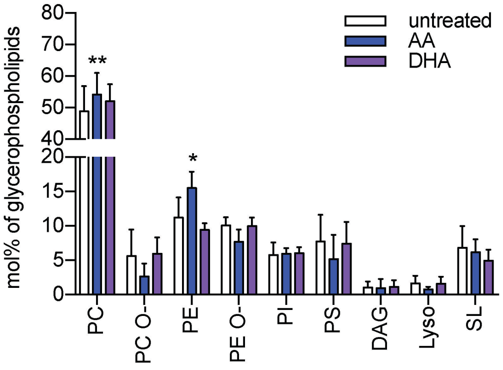


**Fig S5. Minimal changes in lipid head group distribution with PUFA supplementation**. Average ± SD of n ≥ 4 independent experiments. Statistics performed using GraphPad Prism with the two-stage linear step-up procedure of Benjamini, Krieger and Yekutieli, with Q = 1%. Each row was analyzed individually, without assuming a consistent SD. * p < 0.05, ** p < 0.01.

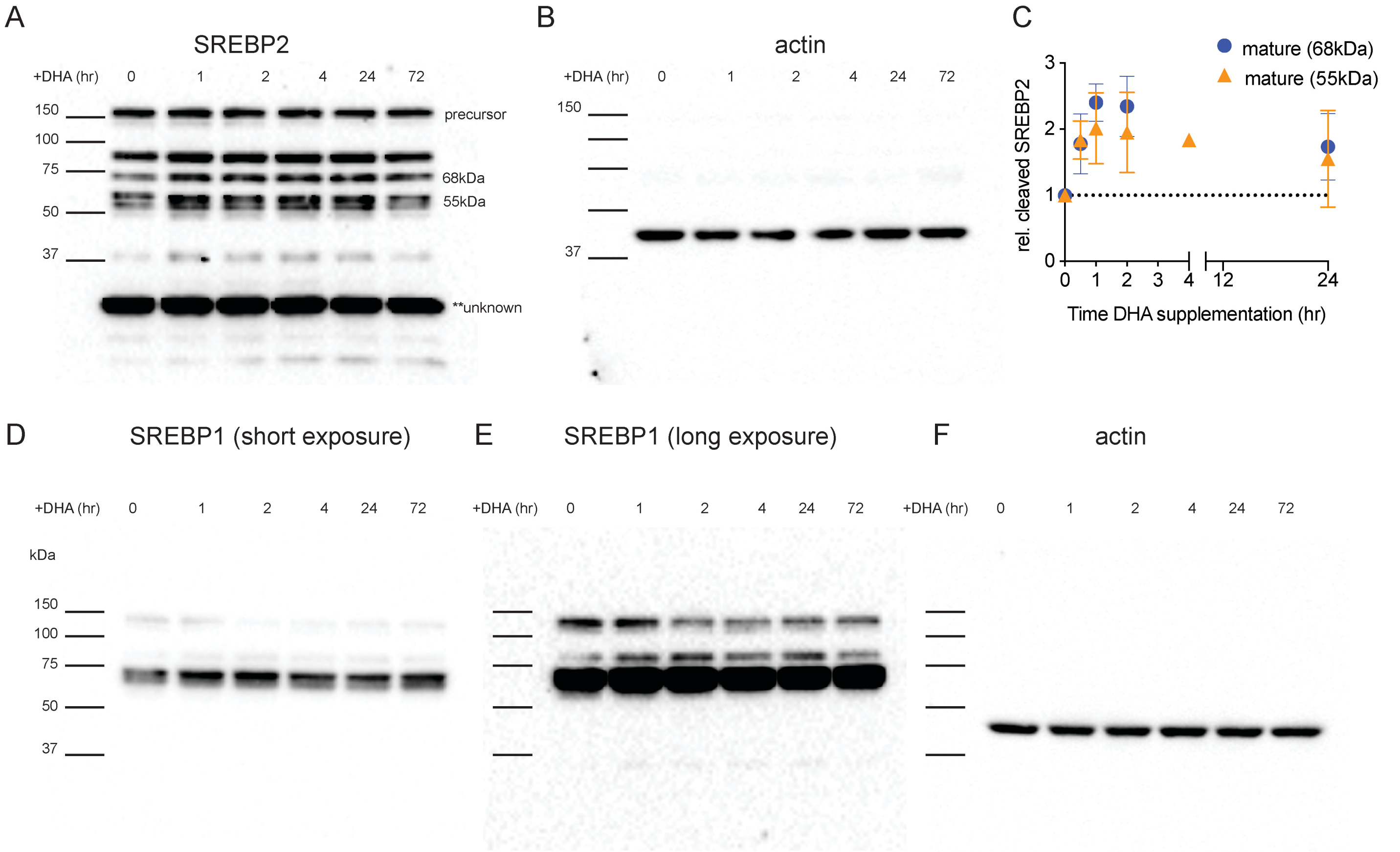


**Fig S6. Full length Western blots for (A-B) SREBP2 and SREBP1 (D-F).** Panel A indicates which bands were quantified in this paper. Panel B shows the same blot reprobed for actin. All Western blots in this paper show the quantification of the protein of interest normalized to actin on the same blot. As there are quite a few unknown bands with this SREBP2 antibody, we quantified two different bands (55 and 68kDa) which run at the approximate size of the cleaved, mature SREBP2 protein. Panel C shows the comparison of the quantification of these two different bands, illustrating that the trend is the same regardless of which band is quantified. Panels D and E show the same Western blot probing for SREBP1 imaged at short (D) or long exposure (E) times. Panel F is the same blot reprobed for actin. These results are representative for n=4-6 individual experiments for SREBP2; n=3 individual experiments for SREBP1.

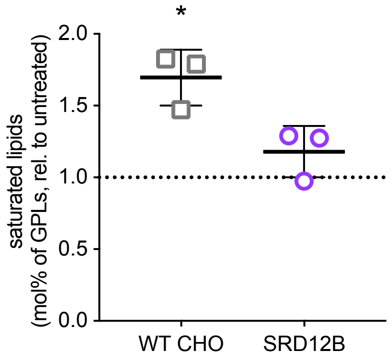


**Fig S7. Lack of saturated lipid upregulation in SRD-12B cells.** Saturated lipids are significantly increased in wild-type, but not SRD-12B, CHO cells treated with 20 µM DHA for 72 h. Shown is the relative mol% of saturated glycerophospholipids compared to untreated cells. Average ± SD of n=3 independent experiments. * $p<0.05$ for one-sample t-test.

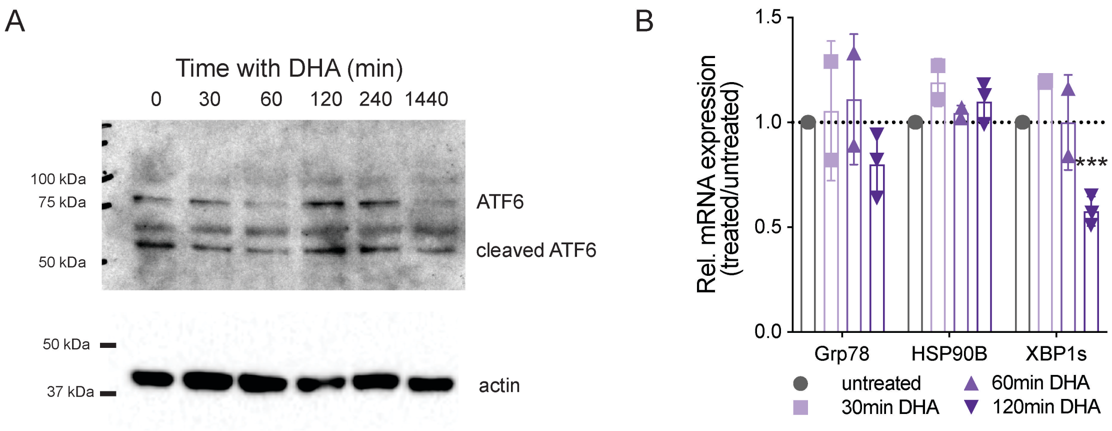


**Fig S8. Proteolytic cleavage of ATF6 is not notably affected by DHA supplementation.** (A) Supplementation of RBL cells with 20 µM DHA does not induce the cleavage of ATF6 (Bethyl Labs, Rabbit polyclonal antibody) as seen by Western blotting for whole cell lysates. Actin is shown as a loading control. (B) qPCR analysis reveals that expression of genes induced by the unfolded protein response is not increased by DHA supplementation, suggesting UPR is not involved in the initial homeostatic response.

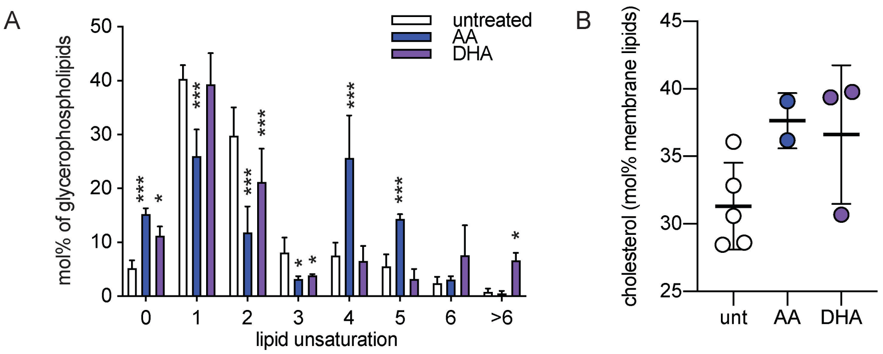


**Fig S9. Lipidomic remodeling in plasma membranes.** (A) Giant Plasma Membrane Vesicles (GPMVs) were isolated from cells treated with 20 µM AA or DHA for 3 days. Lipidomics analysis performed on these isolated PMs shows a similar remodeling response as in the whole cell lipidomes (compare Fig 2). The DHA data set was described in our previous publication[2]. (B) Cholesterol abundance in PMs is increased after three days of supplementation with AA or DHA compared to untreated cells.

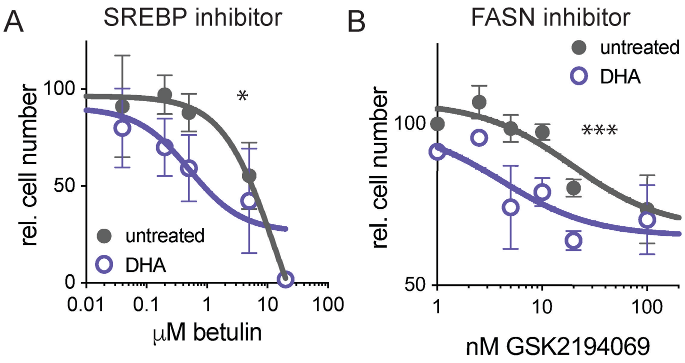


**Fig S10. Dose-response curves for cytotoxic effects of nodes in lipid and/or cholesterol synthesis in the presence of DHA supplementation.** All experiments in this manuscript are performed with 20 µM DHA supplementation, which alone causes no cytotoxicity. Toxicity of inhibiting (A) SREBP maturation (betulin) or (B) fatty acid synthase (GSK2194069) was significantly enhanced by perturbing the lipidome with DHA. Average ± SD of $n \geq 3$ independent experiments. ** $p < 0.05$, *** $p < 0.001$; two-way ANOVA for effect of treatment with 20 µM DHA.

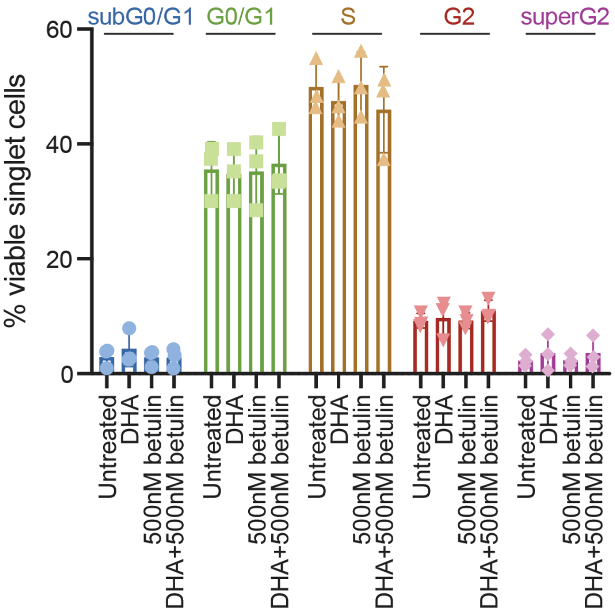


**Fig S11. Cell cycle analysis.** Cell cycle analysis via Hoechst 33342 of DHA- and/or betulin-treated RBL cells shows no difference in cell cycle population with the various treatments. Cells were treated for 24 hr prior to flow cytometry analysis. Average ± SD of n = 3 independent experiments.

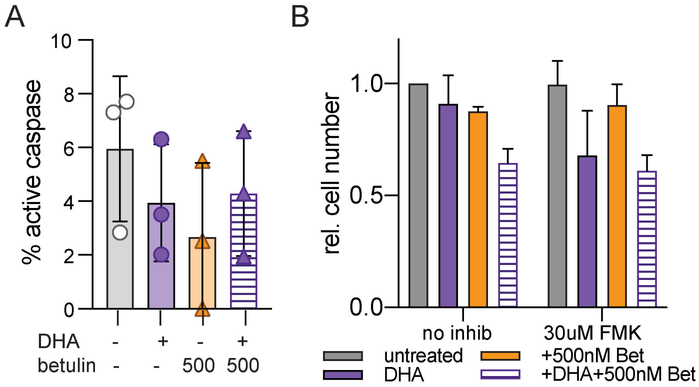


**Fig S12. Cytotoxicity resulting from inhibited membrane homeostasis is not via apoptosis.** (A) RBL cells were treated with 20 µM DHA and/or 500 nM betulin for 24 hr and stained for active Caspase 3/7 (see Materials and Methods). Shown is the percentage of cells positive for active Caspase 3/7, with no significant differences between the four groups. (B) Cells were treated with 20 µM DHA and/or 500nM betulin for 72 hr +/- caspase inhibitor Z-DEVD-FMK. There were no significant differences induced by FMK, suggesting that apoptosis was not involved in cell death induced by combined DHA+betulin treatment.

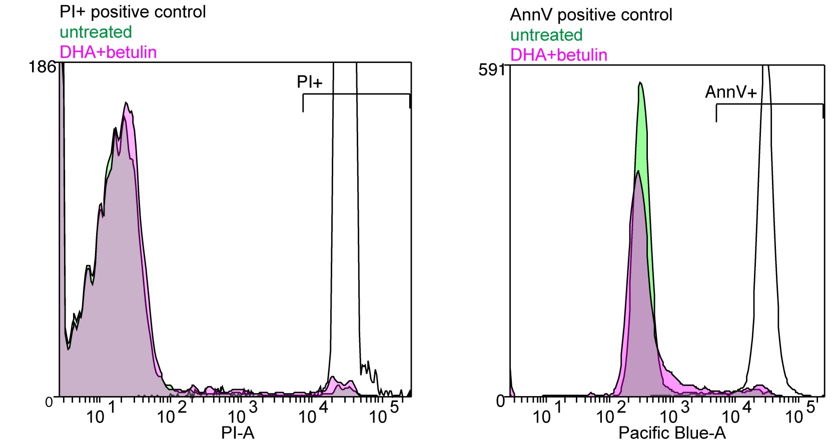


**Fig S13. Histogram of positive controls to establish thresholds for PI or AnxV staining in Fig 6D-G.**

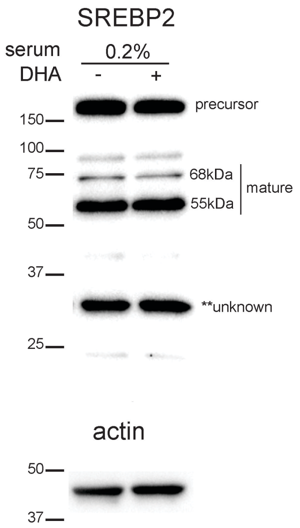


**Fig S14 – Effect of DHA on SREBP2 processing in reduced serum.** RBLs grown in reduced (0.2%) serum overnight with or without 20 μM DHA. In low serum conditions, DHA supplementation does not stimulate SREBP2 proteolysis as shown previously[3], in contrast to full serum (see Fig 4D and Fig S5). Shown is a representative Western blot of SREBP2 and actin (as loading control).

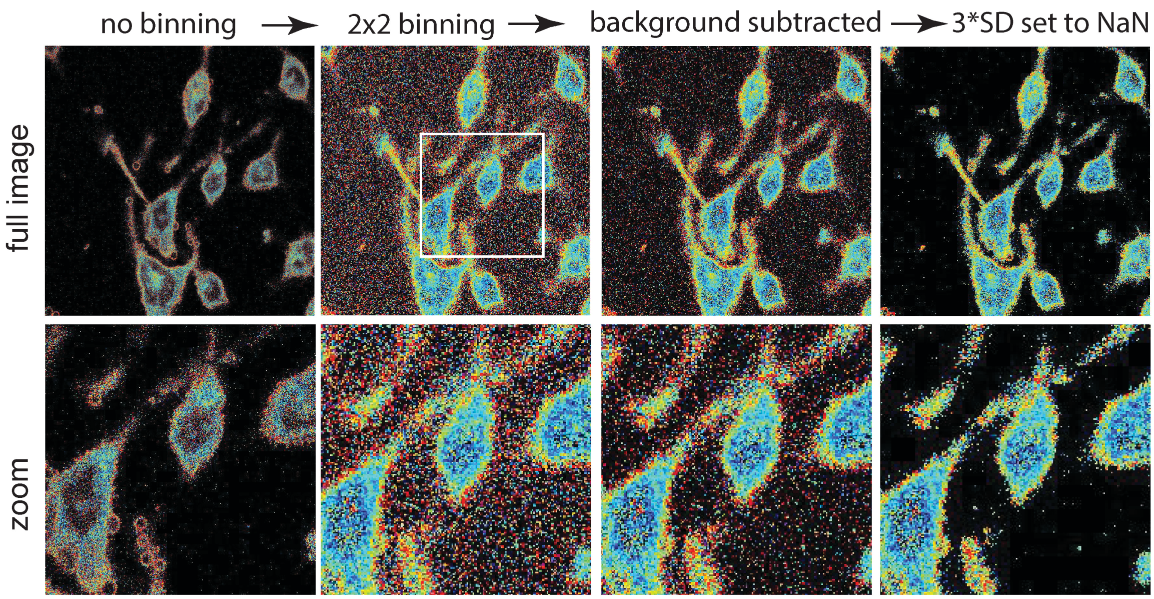


**Fig S15 - Laurdan quantification workflow**. The raw GP map is shown in the left panels. Black pixels are those where GP is undefined due to no signal in either channel. The first step is summative 2x2 binning, which smoothens the imags, but increases noise in the background (adding 4 pixels leads to less zero-containing pixels). Background subtraction is done by selecting a square of pixels outside of the cells and taking an average GP therein. Finally, the standard deviation of those background pixels is calculated and any pixel within 3SD of zero is eliminated from the calculation (NaN rather than zero, because GP=0 is a defined value). Analysis of the rightmost panels shows that minimal data from the cells is eliminated by this procedure.

**Dietary Fatty Acid Composition** [a, b]

| fatty acid [c] | CO [d] | FO [d] |
|---|---|---|
| 14:0 | tr | 7.9 |
| 16:0 | 11.2 | 16.9 |
| 16:1 (n-7) | tr | 10.0 |
| 18:0 | 2.1 | 2.9 |
| 18:1 (n-9) | 30.0 | 13.6 |
| 18:2 (n-6) | 53.8 | 13.1 |
| 18:3 (n-3) | 1.3 | 1.3 |
| 20:5 (n-3) | tr | 11.6 |
| 22:5 (n-3) | tr | 1.6 |
| 22:6 (n-3) | tr | 8.0 |

[a] adapted from ref 3
[b] abbreviations used:
CO - 5 g/100 g corn oil
FO - 4 g/100 g fish oil + 1 g/ 100 g corn oil
[c] only the major fatty acids (>1 g/100 g) are listed
[d] g / 100 g fatty acids

**Table S1.** Fatty acid composition of the corn or fish oil fed to mice for two weeks prior to tissue isolation and lipidomic analysis.


1 Levental, K. R. *et al.* omega-3 polyunsaturated fatty acids direct differentiation of the membrane phenotype in mesenchymal stem cells to potentiate osteogenesis. *Science advances* **3**, eaao1193, (2017).

2 Levental, K. R. *et al.* Polyunsaturated lipids regulate membrane domain stability by tuning membrane order. *Biophys J* **110(8)**, 1800-1810, (2016).

3 Hannah, V. C. *et al.* Unsaturated fatty acids down-regulate srebp isoforms 1a and 1c by two mechanisms in HEK-293 cells. *J Biol Chem* **276**, 4365-4372, (2001).
